# The orphan ligand, Activin C, signals through activin receptor-like kinase 7

**DOI:** 10.1101/2022.03.16.484571

**Authors:** Erich J. Goebel, Luisina Ongaro, Emily Kappes, Elitza Belcheva, Roselyne Castonguay, Ravindra Kumar, Daniel J Bernard, Thomas B. Thompson

## Abstract

Activin ligands are formed from two disulfide-linked inhibin β subunit chains. They exist as homodimeric proteins, as in the case of activin A (ActA; InhβA/InhβA) or activin C (ActC; InhβC/InhβC), or as heterodimers, as with activin AC (ActAC; InhβA:InhβC). While the biological functions of ActA and activin B (ActB) have been well-characterized, little is known about the biological function of ActC or ActAC. One thought is that the InhβC chain functions to interfere with ActA production by forming less active ActAC heterodimers. Here, we assessed and characterized the signaling capacity of ligands containing the InhβC chain. ActC and ActAC activated SMAD2/3-dependent signaling via the type I receptor, activin receptor-like kinase 7 (ALK7). Relative to ActA and ActB, ActC exhibited lower affinity for the cognate activin type II receptors and was resistant to neutralization by the extracellular antagonist, follistatin. In mature adipocytes, which exhibit high ALK7 expression, ActC elicited a SMAD2/3 response similar to ActB, which can also signal via ALK7. Collectively, these results establish that ActC and ActAC are active ligands that exhibit a distinct signaling receptor and antagonist profile compared to other activins.

## Introduction

The activins are multifunctional secreted proteins that play critical roles in growth, differentiation, and homeostasis in a wide variety of cell types. As part of the greater TGFβ family, the activins are dimeric in nature and built from two inhibinβ (Inhβ) chains of approximately 120 amino acids (e.g., activin A, ActA, is built from two InhβA chains) that are tethered by a disulfide bond. Members of the activin class include ActA, activin B (ActB), activin C (ActC), and activin E (ActE), and extend to include GDF8 (myostatin) and GDF11. The Inhβ chains share high sequence identity such that InhβA and InhβB are 63% identical, with InhβC ~50% identical to both InhβA and InhβB^1^. In addition to homodimer formation, several combinations of heterodimers have been observed, such as ActAB formed between InhβA and InhβB chains, as well as the heterodimer ActAC comprised of InhβA and InhβC chains^2–5^. While heterodimers have can form, most studies have focused on the homodimeric forms of the ligands. In addition, due to their established biological roles, many studies have focused on characterizing the ligands ActA and ActB; however, few studies have characterized the ligands ActC or ActE, especially regarding their ability to signal.

The InhβC subunit was first identified from a human liver cDNA library^6^. Its biological role was initially unknown due to the absence of hepatic phenotypes in *Inhbc* knockout mice^7^. Expression of InhβC is highest in the liver but has also been detected in reproductive tissues^8^. InhβC has been proposed to function as an ActA antagonist, as coexpression of InhβA and InhβC results in the formation of the heterodimer ActAC, which is a less active signaling molecule than ActA^4,8^. For example, ActAC is less potent than ActA in IH-1 myeloma cells^9^. Thus, inhβC expression in the presence of InhβA not only reduces ActA levels, but also forms the less potent ligand ActAC. It has also been proposed that ActC directly antagonizes ActA signaling (ref). The mechanism for this is thought to be binding of a non-signaling ActC to the ActA receptors, acting as a competitive inhibitor. These two mechanisms are similar to how inhibin-α (Inhα) forms heterodimers with InhβA chains to form inhibin A, reducing ActA production, and by competitively blocking ActA receptor binding^10^. The similarities were confirmed through studies which showed that in Inhα knockout mice, which develop female reproductive tumors and have abnormally high levels of ActA resulting in cachexia, the ActA levels can be suppressed by over-expression of InhβC^11,12^. While almost all studies have suggested an inhibitory role of InhβC, a recent study showed that ActC relieved chronic neuropathic pain in mice and rats, functioning similarly to TGFβ1, suggesting an agonistic role of the ActC ligand^13,14^.

TGFβ ligands are processed from precursor proteins comprised of a pro-domain, which aids in proper folding and ligand maturation, and a C-terminal signaling domain, which forms the covalent dimers (Fig. 1A). The latter assemble receptor complexes on the cell surface containing a symmetrical positioning of 2 type I and 2 type II serine-threonine kinase receptors, which results in the activation of a SMAD signaling cascade (Fig. 1B). There are seven type I receptors in the family termed activin receptor-like kinases 1 through 7 (ALK1-7)^15^. For ligands of the activin class, the type II receptors bind with high-affinity (nM) to each individual chain in the dimer, with the low-affinity type I receptors binding at a composite interface formed by the two dimer chains (Fig. 1A)^16,17^.

**Figure 1.**
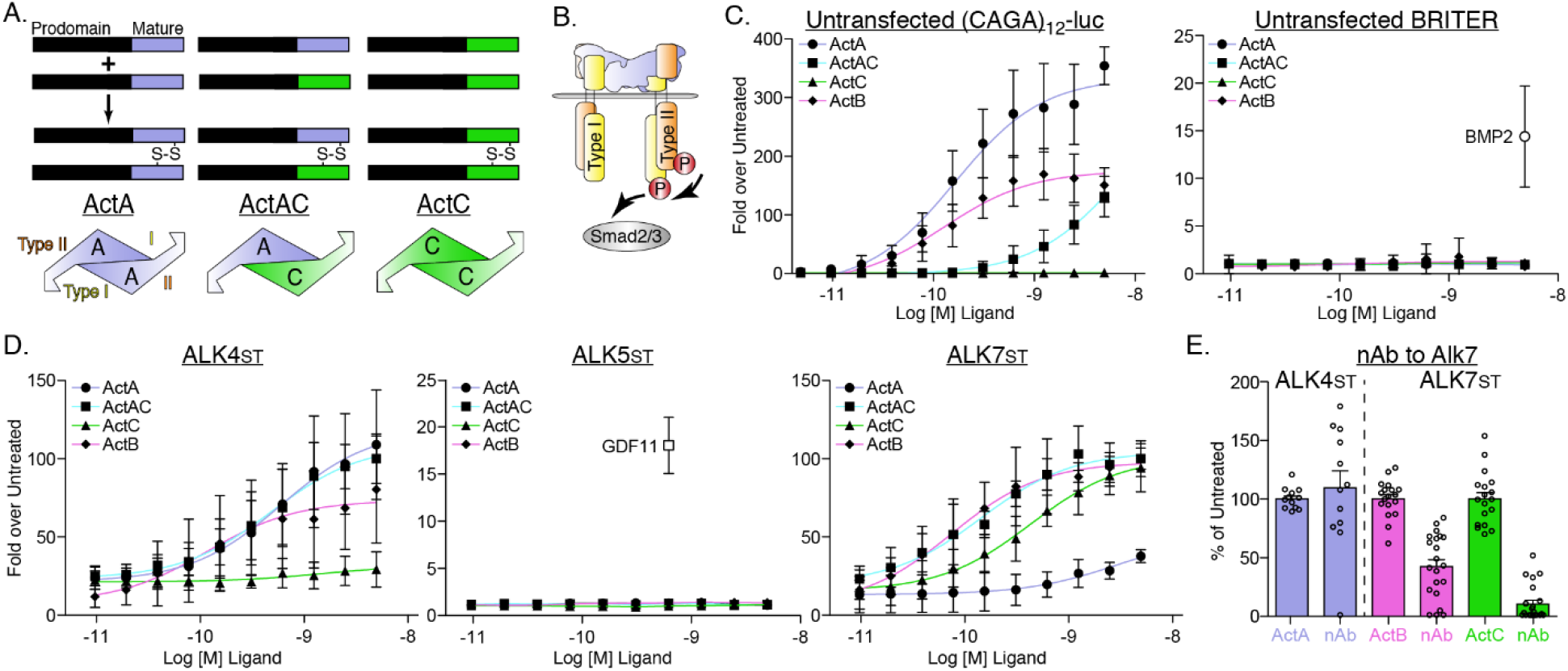
Differences in type I receptor utilization by ActA, ActAC, and ActC. (*A*) Schematic displaying formation of activin A (ActA), AC (ActAC), and C (ActC) from dimerization of inhibin beta A (*blue*) and beta C subunits (*green*). (*B*) Generalized TGFβ signaling schematic displaying activin-SMAD2/3 signaling with type II (*orange*) and type I (*yellow*) receptor binding positions displayed for ActA. (*C*) Luciferase reporter assay in response to ActA, ActAC, or ActC titration in untransfected (CAGA)_12_-luc or BRITER HEKT cells. BMP2 was included as a positive control for the BRITER reporter. (*D*) ActA, ActAC, ActC, and ActB activation of (CAGA)_12_-luc HEK293T cells transfected with SB-431542-resistant (Ser to Thr, ST) type I receptors. In C and D, each data point represents the mean ± SD of triplicate experiments measuring relative luminescence units (RLU). ALK4_st_ and ALK7_st_ transfection assays in D were normalized to 100 fold from mean of highest point. (*E*) Effects of an ALK7 neutralizing antibody (nAb) on ActA, ActB, and ActC induction of (CAGA)_12_-luc activity in cells expressing the indicated type I receptors. In *E,* each data point represents a technical replicate within triplicate experiments with bars displaying the mean ± SD. In both (D) and (E), cells were treated with 10uM SB-431542 to inhibit signaling activity of endogenous receptors.

The activins, as a class, bind to and signal through three type I receptors: ALK4, ALK5 and ALK7. In general, each member signals through ALK4, whereas GDF8 and GDF11 extend specificity to ALK5 and ActB to ALK7^18,19^. Structural and biochemical studies have made strides in illuminating the determinants of specificity between the activins and the type I receptors, providing context for the different biological roles of each activin member^16,20-22^. ALK4 and ALK5 expression is relatively widespread, while ALK7 is primarily expressed in the adipose and reproductive tissues, and also in brain and pancreatic cells^23,24^. While ALK7 specific signaling has been linked to cancer cell apoptosis, its role in adipose tissue is more well-studied^25,26^. ActB signaling via ALK7 in adipose tissue suppresses lipolysis and downregulates adrenergic receptors, facilitating fat accumulation^27-29^. Similarly, loss of signaling in ALK7 knockout mice renders the animals resistant to diet-induced obesity^27-29^.

Unlike the other activin ligands, limited information is available for the signaling capacity of ActC and whether the homodimer can actually activate SMAD molecules. One hypothesis is that ActC is a non-signaling molecule and simply a non-functional by-product of expressing InhβC in the presence of InhβA. Given this uncertainty, we sought to characterize the signaling capacity of ActC across the panel TGF-β type I receptors. We demonstrate that homodimeric ActC can act as a potent activator of SMAD2/3 and does so with high specificity via the type I receptor, ALK7. Additionally, unlike the rest of the activin class, ActC has a much lower affinity for the type II receptors, ActRIIA and ActRIIB. Intriguingly, ActC is not antagonized by follistatin, which potently neutralizes ActA, ActB, GDF8, and GDF11. Finally, we demonstrate that ActC can activate SMAD2/3 signaling similarly to ActB in mature adipocytes in an ALK7-dependent manner.

## Methods

### Protein expression and purification

#### ActRIIA and ActRIIB

The extracellular domains of human ActRIIA (residues 1-134) and rat ActRIIB (residues 1-120) were produced as previously described^16^. Specifically, both receptors were subcloned into the pVL1392 baculovirus vector with C-terminal Flag and His_10_ tags (ActRIIA) or a C-terminal His_6_ tag followed by a thrombin cleavage site (ActRIIB). Recombinant baculoviruses were generated through the Bac-to-Bac system (ActRIIA; Invitrogen - Waltham, MA) or the Baculogold system (ActRIIB; Pharmingen - San Diego,CA). Virus amplification and protein expression were carried out using standard protocols in SF+ insect cells (Protein Sciences - Meriden, CT). ActRIIA and ActRIIB were purified from cell supernatants by using Ni Sepharose affinity resin (Cytiva – Marlborough, MA) with buffers containing 50mM Na_2_HPO_4_, 500mM NaCl, and 20mM imidazole, pH 7.5 for loading/washing and 500mM imidazole for elution. ActRIIB was digested with thrombin overnight to remove the His_6_ tag. ActRIIA and ActRIIB were subjected to size exclusion chromatography (SEC) using a HiLoad Superdex S75 16/60 column (Cytiva) in 20mM Hepes, and 500mM NaCl, pH 7.5.

#### ActRIIA-Fc, ActRIIB-Fc, ActRIIB-ALK7-Fc, ALK7-Fc and ActRIIB-ALK4-Fc

ActRIIA-Fc was purchased from R&D (Cat. No. 340-RC2-100 - Minneapolis, MN). ActRIIB-Fc was expressed and purified from Chinese hamster ovary cells as previously described^21,56^. Briefly, ActRIIB-Fc was isolated using affinity chromatography with Mab Select Sure Protein A (GE healthcare - Waukesha, WI), followed by dialysis into 10mM Tris, 137mM NaCl, and 2.7mM KCl, pH 7.2. ActRIIB-ALK7-Fc and ActRIIB-ALK4-Fc were designed and expressed as previously described in CHO DUKX cells through the coexpression of two plasmids, each containing a receptor ECD (ActRIIB or ALK4) fused to a modified human IgG1 Fc domain^34,57^. Purification was performed through protein A MabSelect SuRe chromatography (Cytiva), then eluted with glycine at low pH. The resulting sample was further purified over a Ni Sepharose 6 fast flow column (Cytiva) followed by an imidazole elution gradient, an ActRIIB affinity column and ultimately, a Q Sepharose column (Cytiva). ALK7-Fc production was performed as described previously^57^.

#### Antibodies neutralizing ActRIIA, ActRIIB, ActRIIA/ActRIIB and ALK7

An Anti-ActRIIA antibody was obtained through phage-display technology, while anti-ActRIIB, anti-ActRIIA/ActRIIB and ALK7 antibodies were generated using the Adimab platform. The first three antibodies were expressed from stable CHO pools, while anti-ALK7 was transiently expressed in ExpiCHO cells (Thermo Fisher). The CM was purified over Mab SelectSure Protein A (Cytiva) followed by ion-exchange chromatography.

#### ActA, ActB, and Inhibin A

Mature, recombinant ActA and ActB were prepared as previously described^16,19^. Briefly, ActA (pAID4T) was expressed in Chinese hamster ovary (CHO) DUKX cells. Conditioned media (CM) of ActA was then mixed with a proprietary affinity resin made with an ActRIIA-related construct (Acceleron). The resin was then lowered to pH 3 to dissociate the propeptide-ligand complex. Following this, the pH was raised to 7.5 and the resin was incubated for 2h at room temperature. ActA was eluted with 0.1M glycine pH 3.0, which was concentrated over a phenyl hydrophobic interaction column (Cytiva) and eluted with 50% acetonitrile/water with 0.1% trifluoroacetic acid (TFA). Lastly, ActA was further purified by HLPC over a reverse phase C4 column (Vydac) with a gradient of water/0.1% TFA and acetonitrile/0.1% TFA. Expression of ActB was performed through the use of a previously generated CHO-DG44 stable cell line^19^. CM was initially clarified over an ion exchange SP XL column (Cytiva) in 6M urea, 25mM MES, 50mM Tris pH 6.5. The flow-through was then adjusted to 0.8M NaCl and applied to a Phenyl Sepharose column (Cytiva) and ActB was eluted by decreasing the NaCl through a gradient. Lastly, ActB was purified by reverse phase chromatography (C18, Cytiva) and eluted similarly to ActA. Recombinant human ActB used in the assays involving differentiated adipocytes was purchased from R&D (Cat. No. 659-AB-005). Inhibin A was produced and purified as previously described^58^.

#### ActAC and ActC

Mature recombinant human ActAC and ActC were purchased from R&D (Cat. No. 4879-AC and 1629-AC, respectively). Antibodies used include: ActA (AF338, R&D); ActC (MAB1639, R&D); Goat (PI-9500, Vector Laboratories) and Mouse (DC02l, Calbiochem). ActC wt (IQP) and ActC Gln267Ala (IAP) were expressed transiently in HEK293T cells using a construct with an optimized furin cleavage site. The conditioned media was then adjusted to 0.8M NaCl and applied to a Phenyl Sepharose column (Cytiva) followed by elution with low NaCl. Lastly, ActC was then purified using reverse phase chromatography (C18, Cytiva) and eluted similarly to ActA and ActB.

#### Fst288 and Fstl3

Both Fst288 and Fstl3 were produced as previously described^39^. Fst288 was expressed from a stably transfected CHO cell line and purified from CM by binding to a heparin-sepharose column (abcam) in 100mM NaBic pH 8 and 1.5M NaCl, with a low salt gradient to elute followed by cation exchange over a Sepharose fast flow (Cytiva) in 25mM HEPES pH 6.5, 150mM NaCl with a high salt elution gradient. Finally, Fst288 was then purified over an HPLC SCX column in 2.4 mM Tris, 1.5 mM Imidazole, 11.6 mM piperazine pH 6 with a high salt, high pH (10.5) gradient elution. Fstl3 was cloned into the pcDNA3.1/myc-His expression vector and expressed transiently in HEK293F cells. CM was harvested after 6 days and applied to His-affinity resin (Cytiva), followed by washing with a buffer of 500mM NaCl, 20mM Tris pH 8 and elution with 500mM imidazole. Fstl3 was then subjected to SEC using a HiLoad Superdex S75 16/60 column (Cytiva) in 20mM HEPES pH 7.5 and 500mM NaCl.

### Luciferase reporter assays

Assays using the HEK-293-(CAGA)_12_ or BRITER luciferase reporter cells were performed in a similar manner as described previously^16,19,20,59^. Specifically, cells were plated in a 96-well format (3 x 10^4^ cells/well) and grown for 24h. For standard EC50 experiments (Fig. 1B and C), growth media was removed and replaced with serum free media supplemented with 0.1% BSA (SF^BSA^ media, Thermo Fisher) and the desired ligand, where a two-fold serial dilution was performed with a starting concentration of 160nM (ActA and ActAC, (CAGA)_12_) or 4.96nM (ActC, (CAGA)_12_ and ActA, ActAC, ActC, BRITER). Incubation was performed for 18 h, cells were then lysed and assayed for luminescence using a Synergy H1 Hybrid plate reader (BioTek – Winooski, VT). For the assays featuring transfections of ALK4_st_, ALK5_st_ or ALK7_st_, a total of 50ng DNA (10ng Type I receptor, 40ng empty vector) was transfected using Mirus LT-1 transfection reagent at 24h post-plating. Each receptor construct contains a single point mutation (pRK5 rat ALK5 S278T (ST), pcDNA3 rat ALK4 S282T, pcDNA4B human ALK7 S270T) conferring resistance to the inhibitory effects of the small molecule SB-431542. Media was then removed and replaced with SF^BSA^ media with 10μM SB-431542 and the desired ligand for 18h. For the experiments featuring ActRIIA-Fc, ActRIIB-Fc, the neutralizing antibodies, Fst288 or Fstl3, these proteins were added to the ligands and incubated for 10min prior to addition to cells. The luminescence data was imported into Graphpad Prism for figure generation.

### Surface plasmon resonance (SPR) studies

SPR experiments were carried out in HBS-EP+ buffer (10mM HEPES pH 7.4, 500mM NaCl, 3.4 mM EDTA, 0.05% P-20 surfactant, 0.5mg/ml BSA) at 25C on a Biacore T200 optical biosensor system (Cytiva). Fc-fusion constructs of each receptor were captured using either a Series S Protein A sensor chip (GE Healthcare) or a Series S CM5 sensor chip (GE Healthcare) with goat anti-human Fc-specific IgG (Sigma-Aldrich – Saint Louis, MO) immobilized with a target capture level of ~70 RU. Experiments with ActRIIA-Fc (ActA, ActAC, ActB), ActRIIB-Fc (ActA, ActAC, ActB), and ActRIIB-ALK4-Fc (ActA, ActAC, ActC) were performed with the former chip while experiments coupling ActRIIA-Fc (ActC), ActRIIB-Fc (ActC) and ActRIIB-ALK7-Fc (ActA, ActAC, ActC, ActB) were performed with the latter chip methodology. An 8-step, two-fold serial dilution was performed in the aforementioned buffer for each ligand, with an initial concentration of 10nM (for Activin C, a 10-step, two-fold serial dilution beginning at 150 nM was performed, for Activin AC, a 9-step, two-fold serial dilution starting at 20nM was performed). Each cycle had a ligand association and dissociation time of 300 and 600 seconds, respectively. The flow rate for kinetics was maintained at 50uL/min. SPR chips were regenerated with 10mM Glycine pH 1.7. Kinetic analysis was conducted using the Biacore T200 evaluation software using a 1:1 fit model with mass transport limitations (red lines). Each binding experiment was performed in duplicate, fit individually and then averaged.

### Structural modeling and alignments

The model of ActC was built with Swiss-model using several ActA structures as templates: PDB codes 1S4Y (ActA:ActRIIB), 2ARV (unbound ActA), 2B0U (ActA:Fs288), 5HLZ (Pro-ActA) and 7OLY (ActA:ActRIIB:ALK4)^20,38,60-63^. A consensus was observed in the overall structure, particularly at the type II interface and IAP motif. Ultimately, the model built from 7OLY was used for the comparison in Figure 5, as it is the most complete ActA-receptor complex, and all images and alignments were performed in PyMol (The PyMol Molecular Graphics System, Schrödinger, LLC, New York, NY).

### Adipocyte isolation, differentiation, and treatment for western blot

Adipocyte stem cells were isolated, cultured, and differentiated as previously described^64^. Briefly, inguinal adipose tissue was harvested aseptically from male mice (3-4 weeks old) and placed in sterile PBS, followed by mincing and collagenase digestion (1 mg/ml) for 1 h at 37°C. Then, the digestion was filtered through a 70-μm mesh and centrifuged to separate the stromal vascular fraction (SVF). Following aspiration, the SVF was resuspended in DMEM supplemented with 10% FBS and Pen-strep-amphotericin (Wisent Inc. cat. No: 450-115-EL - Saint-Jean-Baptiste, Canada) and plated in a 6-well format at ~320,000 cells/well. Following expansion over four days, cells were differentiated over the course of four days using a solution of 5μM dexamethasone, 0.5mM 3-isobutyl-1-methylxanthine, 10μg/ml insulin and 5μM Rosiglitazone. Adipocytes were then maintained for six additional days prior to experimentation in DMEM/FBS + insulin. 3T3-L1 cells were differentiated to adipocytes following ATCC recommended protocol. Briefly, 3T3-L1 cells were differentiated over the course of four days using a solution of 1 μM dexamethasone, 0.5 mM 3-isobutyl-1-methylxanthine and 1 μg/ml insulin. 3T3-L1 chemically-induced adipocytes were then maintained for six additional days prior to experimentation in DMEM/FBS + insulin. Differentiated adipocytes from SVF or 3T3-L1 cells were then starved in serum-free media for 1h, after which they were treated with serum-free media containing ActA, ActB or ActC (2nM) for 1h + Fst288 (800 ng/ml). In another set of experiments, differentiated-SVF cells were treated with ActA, ActB or ActC (2nM) for 1h + anti-ALK7 antibody (30 μg/ml). Concentrations were selected based on in vitro cell-based assays. At the end of the treatments, cells were lysed using RIPA buffer containing protease inhibitors and western analysis was performed using anti-pSmad2 (Cell Signaling, 138D4 - Danvers, MA) or anti-SMAD2/3 antibodies (Millipore, 07-408 - Burlington, MA).

### Adipocyte RNA extraction

Cells were collected in TRIzol and RNA was extracted following the manufacturer’s protocol (Zymo Research). Total RNA from SVF or 3T3-L1 adipocyte-differentiated cells (at day 10 of differentiation) (200 ng) was reverse transcribed using (MMLV) reverse transcriptase following the manufacturer’s protocol (Promega - Madison, WI). Expression of genes encoding the *Pparγ2, Cebpa*, and *Pnpla2* was analyzed in duplicate qPCR reactions using EvaGreen Master mix (ABMMmix-S-XL; Diamed) on a Corbett Rotorgene 6000 instrument (Corbett Life Science, Mortlake, NSW, Australia). Gene expression was determined relative to the housekeeping gene *Rpl19* using the 2-ΔΔCt method^65^. Primer sequences are listed in Table S2.

### Adipocyte images and Oil Red O staining

Before and after day 10 of differentiation, adipocyte images were acquired with an Axiocam 506 mono camera (Zeiss - White Plains, NY) using ZEN 2.3 pro (Zeiss) software. For Oil Red O (ORO) staining, cells were washed in PBS and fixed in 10% formalin buffered solution for 10 min. After fixation, cells were washed in 60% isopropanol and stained in an ORO solution (2:3 v/v H_2_O: isopropanol, containing 0.5% ORO, Sigma O0625) for 1 hour. After staining, cells were washed in PBS and dye from lipid droplets was extracted by adding pure isopropanol for 10 min in a rotor shaker. Dye per well was quantified by absorbance at 500 nm in EZ Read 2000 microplate reader (Biochrom - Holliston, MA). After washing with PBS, cells were digested using a 0.25% Trypsin solution in PBS-EDTA for 24 h at 37°C. DNA was quantified using a Nanodrop, and cell lipid content was normalized by the corresponding cell DNA content per well.

## Results

### Activin C induces SMAD2/3 phosphorylation through ALK7

Cell-based reporter assays have long been used to measure SMAD activation. To investigate ActC’s ability to induce canonical SMAD2/3 signaling like other activins, we performed luciferase reporter assays in an activin-responsive HEK293T cell line stably transfected with (CAGA)_12_-luciferase plasmid^19,20^. Purified recombinant activin ligands (ActA, ActAC, ActC, and ActB) were titrated to generate EC50 curves. In this format, ActA stimulated a response at lower ligand concentration than either ActAC or ActB (Fig. 1C, left panel). In contrast, ActC did not induce reporter activity up to concentrations of 5 nM. Of note, ActAC showed about half of the activity of ActA, consistent with ActAC being less potent than ActA but more potent than ActC. Neither ActAC nor ActC activated a SMAD1/5/8-dependent reporter in an osteoblast cell line, in contrast to the robust response observed with BMP2 (Fig. 1C, right panel).

Though the above data show that ActC does not signal like other activin ligands, HEK293 cells endogenously express only two of the three SMAD2/3 type I receptors, ALK4 and ALK5, with little to no expression of ALK7^19,30^. To address this limitation, we applied a heterologous system developed to interrogate specific signaling from individual type I receptors^16^. Here, a point mutation was introduced into each type I receptor [ALK4 S282T (ALK4_st_), ALK5 S278T (ALK5_st_), or ALK7 S270T (ALK7_st_)] that rendered it resistant to the small molecule kinase inhibitor, SB-431542, while maintaining ligand-induced activation. HEK293T (CAGA)_12_-luciferase reporter cells were transiently transfected with the modified receptors, then co-treated with the indicated ligands and SB-431542 to suppress signaling from endogenous receptors. In the presence of ALK4_st_, ActA, ActAC, and ActB, but not ActC, stimulated reporter activity (Fig. 1D). In ALK5_st_-transfected cells, none of the activins stimulated reporter activity, while GDF11, a known ALK5 ligand, served as a positive control (Fig. 1D). As expected, ActB and to a much lesser extent, ActA, induced reporter activation when ALK7_st_ was transfected into the HEK293T (CAGA)_12_-luciferase reporter cells. Strikingly and unexpectedly, ActAC and ActC activated (CAGA)_12_-luc activity in the presence of ALK7_st_ to a similar extent as ActB (Fig. 1D).

These data suggested that ActC can signal specifically through ALK7. To further validate these results, we utilized an antibody that was developed to specifically bind and neutralize ligand signaling through ALK7. Following treatment with the anti-ALK7 antibody, ActB signaling via ALK7st was significantly reduced while ActC signaling was nearly abrogated completely (Fig. 1E). The specificity of the antibody for ALK7 was confirmed as ActA signaling via ALK4_st_ was unaffected (Fig. 1E). Taken together, these data show that the activin ligands have differential type I specificities. ActA signals predominantly through ALK4, ActB and ActAC signal through ALK4 and ALK7, while ActC signals exclusively through ALK7.

### Activin C and Activin AC interact with and require activin type II receptors to signal

In addition to the type I receptors, TGFβ family ligands must bind type II receptors to generate intracellular signals. Ligands of the activin class generally bind the type II receptors ActRIIA and ActRIIB with high affinity (pM-nM)^16,21,31–33^. Previous studies showed that ActAC binds ActRIIB with lower affinity than ActA, suggesting that the InhβC chain has a diminished type II interaction^11^. Given the new finding that ActC is a signaling molecule, we sought to determine and compare the binding affinities of ActA, ActAC, and ActC to ActRIIA and ActRIIB using Surface Plasmon Resonance (SPR). In this experiment, type II receptor extracellular domains (ECDs) fused to an antibody Fc fragment were captured using a Protein A biosensor chip, while ActA, ActAC, or ActC were titrated as the analyte. For both ActRIIA-Fc and ActRIIB-Fc, binding affinity was highest for ActA (equilibrium constant (apparent K_D_) of 22 pM and 8.1 pM, respectively) (Fig. 2, top row). ActAC also bound ActRIIA and ActRIIB with high affinity, although slightly weaker than ActA (150 pM and 90 pM, respectively), suggesting that the InhβC chain diminishes the overall type II affinity of the dimer (Fig. 2, middle row). ActC binding to ActRIIA and ActRIIB was much weaker, with a significantly faster dissociation rate than either ActA or ActAC (Fig. 2, bottom row). While ActA had no apparent preference for one type II receptor, consistent with previous studies, ActC had a higher affinity for ActRIIA than ActRIIB (Fig. S1A)^16^. Similarly, ActB binding to type II receptors was much stronger then ActC (Fig. S1B), indicating that ActC deviates from other activin class ligands by exhibiting low affinity for type II receptors. To confirm the weak binding of ActC towards type II receptors, we performed a native gel analysis where ActA, ActAC, and ActC were incubated with either ActRIIA or ActRIIB. Both ActA and ActAC, but not ActC, formed a stable complex when incubated with either ActRIIA and ActRIIB (Fig. S2). Collectively, these data indicate that ActC deviates from other activin class ligands by exhibiting low affinity for type II receptors.

**Figure 2.**
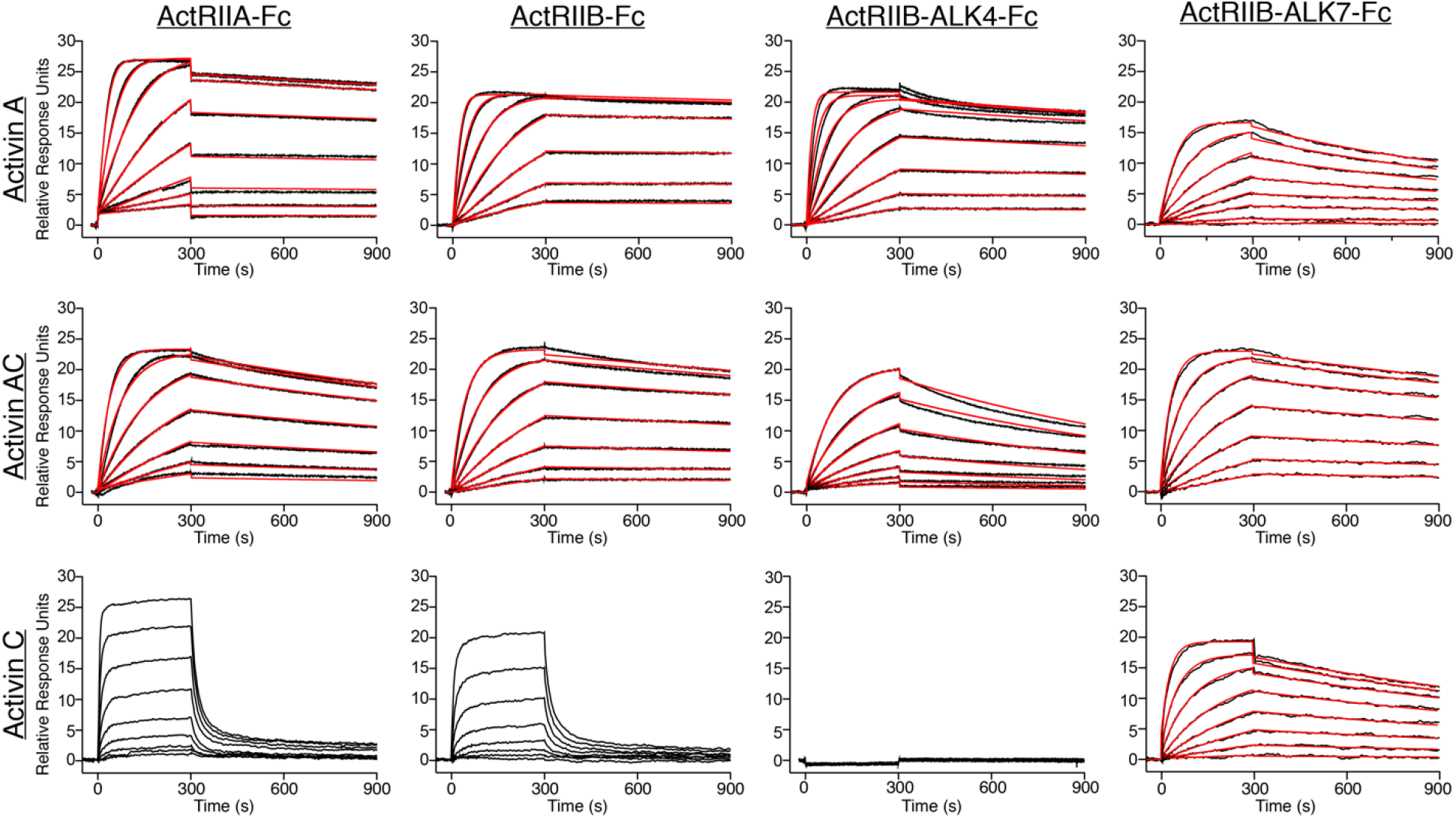
ActC binds activin type II receptors with low affinity. Representative SPR sensorgrams of ActA, ActAC, and ActC binding to protein-A captured ActRIIA-Fc, ActRIIB-Fc, ActRIIB-ALK7-Fc or ActRIIB-ALK4-Fc. Sensorgrams (*black lines*) are overlaid with fits to a 1:1 binding model with mass transport limitations (*red lines*). ActC binding to ActRIIA and ActRIIB were fit using a steady state model. Each experiment was performed in duplicate with the kinetic parameters summarized in SI appendix Table S1.

We also used SPR to determine if ActC or ActAC bind specifically to ALK7. No to little binding of any of the activins, including ActC and ActAC, was observed for ALK7-Fc, indicating that for all ligands ALK7 is a low affinity receptor (Fig. S1C). With a low affinity type II and type I receptor, we asked whether the combination of receptors enhanced binding of ActC. A heterodimeric-Fc receptor fusion that incorporates both the type I and type II receptor can mimic natural signaling pairs^34^. Previously, ActRIIB-ALK4-Fc exhibited higher affinity for ActA than the monovalent ActRIIB-Fc, indicating enhanced binding due to incorporation of the type I receptor^20^. We therefore tested binding of the heterodimeric ActRIIB-ALK7-Fc to ActA, ActAC, and ActC (Fig. 2). ActA bound ActRIIB-ALK7-Fc (296 pM) with 7-fold lower affinity than ActRIIB-Fc (42 pM), indicating ALK7 did not contribute to binding, consistent with ActA not signaling via ALK7. Interestingly, significant binding was observed for both ActC (2 nM) and ActAC (51 pM) to ActRIIB-ALK7. While for ActAC, binding to ActRIIB-ALK7-Fc was slightly higher than binding to ActRIIB-Fc (64 pM), a dramatic difference was observed for ActC where binding was increased 40-fold over ActRIIB-Fc alone. Similar studies were performed with ActRIIB-ALK4-Fc. As expected ActA bound with high-affinity to ActRIIB-ALK4-Fc while ActAC had a much weaker interaction (457 pM), and ActC failed to bind (Fig. 2). SPR experiments with ActB showed consistent results, where high-affinity interactions were observed with ActRIIA, ActRIIB, and ActRIIB-ALK7-Fc, with little to no binding to ALK7-Fc (Fig. S1C). Binding data for each SPR experiment can be found in Table S1.

Next, we investigated whether ActC signaling could be inhibited using the type II receptor-Fc constructs as a competitive antagonist (ligand trap) to block endogenous receptor binding in a cell-based assay (Fig. 3A). We employed the same assay system in which SB-431542 resistant type I receptors were transfected into HEK293T (CAGA)_12_-luc reporter cells. We titrated either ActRIIA-Fc or ActRIIB-Fc against a constant concentration (0.62 nM) of ActA, ActAC, ActC, or ActB (Fig. 3B and C). ActRIIA-Fc and ActRIIB-Fc dose-dependently inhibited ActA and ActAC signaling via ALK4_st_ and ablated ActB signaling via ALK7_st_. In contrast, signaling by both ActAC and ActC through ALK7_st_ was not inhibited by ActRIIA-Fc or ActRIIB-Fc, even at the highest concentration of the decoy receptors (25 nM).

**Figure 3.**
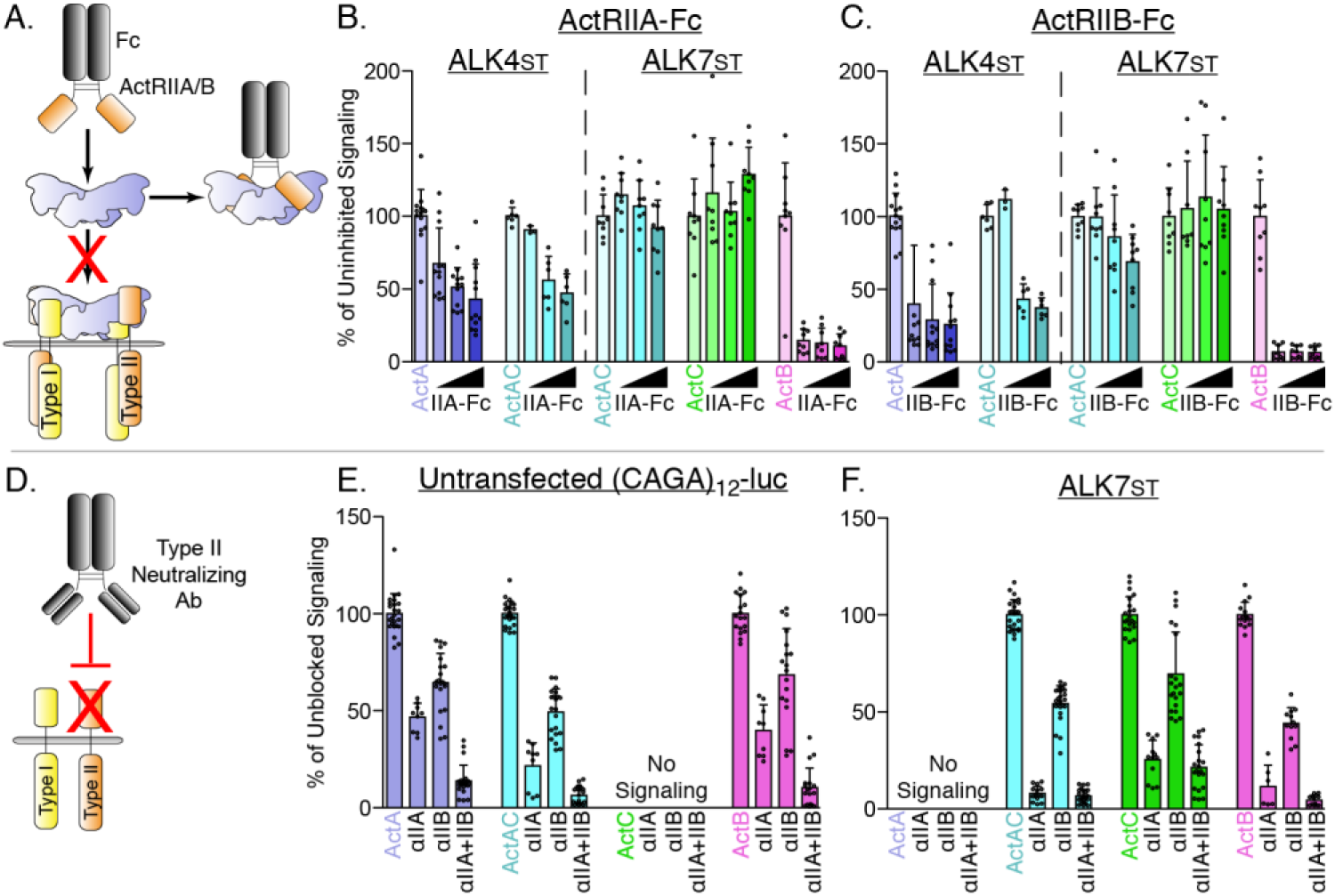
ActA and ActAC signal via activin type II receptors. (*A*) Schematic representation of activin type II receptor Fc-fusion proteins as decoys. (*B and C*) HEK293T (CAGA)_12_-luc cells were transfected with ALK4_st_ and ALK7_st_ and treated with SB-431542 and ActA, ActAC, ActC, or ActB (0.62nM) as in Fig. 1 in the presence of increasing quantities of either ActRIIA-Fc (*B*) or ActRIIB-Fc (*C*). (*D*) Schematic representation of neutralizing antibodies targeting the type II receptor ECDs. (*E and F*) HEK293T (CAGA)_12_-luc cells following treatment with ActA, ActAC, ActC, or ActB (0.62 nM) in the presence or absence of neutralizing antibodies targeting ActRIIA, ActRIIB, or both. No signaling was observed by ActC in *E* or ActA in *F*. Each data point represents technical replicates within triplicate experiments measuring relative luminescence units (RLU) with bars displaying the mean ± SD. Data are represented as % of uninhibited (*B and C*) or unblocked (*E and F*) signal.

Given its low affinity for activin type II receptors, we wanted to determine whether ActC signaling through ALK7 was dependent on ActRIIA or ActRIIB. We therefore used a series of receptor neutralizing antibodies, which bind to either the ECD of ActRIIA or ActRIIB, or to both receptors (Fig. 3D). As expected, neutralization of either ActRIIA or ActRIIB significantly reduced signaling by ActA and ActB in the untransfected and untreated (i.e., without SB-431542) CAGA-luc cells (Fig. 3E). An antibody that simultaneously blocks both ActRIIA and ActRIIB more potently inhibited signaling of each activin ligand than the single-target antibodies, indicating that ActRIIA and ActRIIB were redundant. ActAC signaling in the untransfected/untreated CAGA-luc cells was also inhibited by type II receptor neutralization in a similar manner to ActA and ActB. Again, ActC did not signal under these assay conditions unless ALK7_st_ was added. Here, both ActAC and ActB signaling was readily inhibited by type II receptor blockade (Fig. 3F). ActC signaling was significantly reduced when ActRIIA was blocked and to a lesser extent with blocking ActRIIB. These observations demonstrate that, despite their lower affinities, ActC and ActAC require a type II receptor, with a preference for ActRIIA, for signaling via ALK7.

### Activin C is antagonized by inhibin A, but not follistatin-288 or follistatin-like protein 3

Activin class ligands are regulated through several mechanisms. One such mechanism is through extracellular antagonists, such as follistatin-288 (Fst-288) and follistatin-like 3 (FSTL3), which bind and sequester ligands. Fst-288 and FSTL3 form a donut-like shape to surround activin ligands, occluding epitopes that are important for binding to both type I and type II receptors^35-39^. Given the new finding that ActC can signal via type I (ALK7) and II (ActRIIA/B) receptors, we next examined whether its actions were inhibited by either Fst-288 or FSTL3. Fst288, at two concentrations, robustly inhibited ActA and ActB induction of CAGA-luc activity via ALK4_st_ (Fig. 4A). Fst288 also inhibited ActB signaling via ALK7_st_, though to a lesser extent. Fst288 moderately inhibited ActAC signaling via ALK4_st_ and ALK7_st_, but unexpectedly had no impact on ActC actions (Fig. 4A). Similar results were observed with the related antagonist FSTL3, which binds and occludes activin ligands similarly to Fst288 (Fig. 4B)^39,40^. This unique resistance to classical activin antagonists further distinguishes ActC from the rest of the activin class.

**Figure 4.**
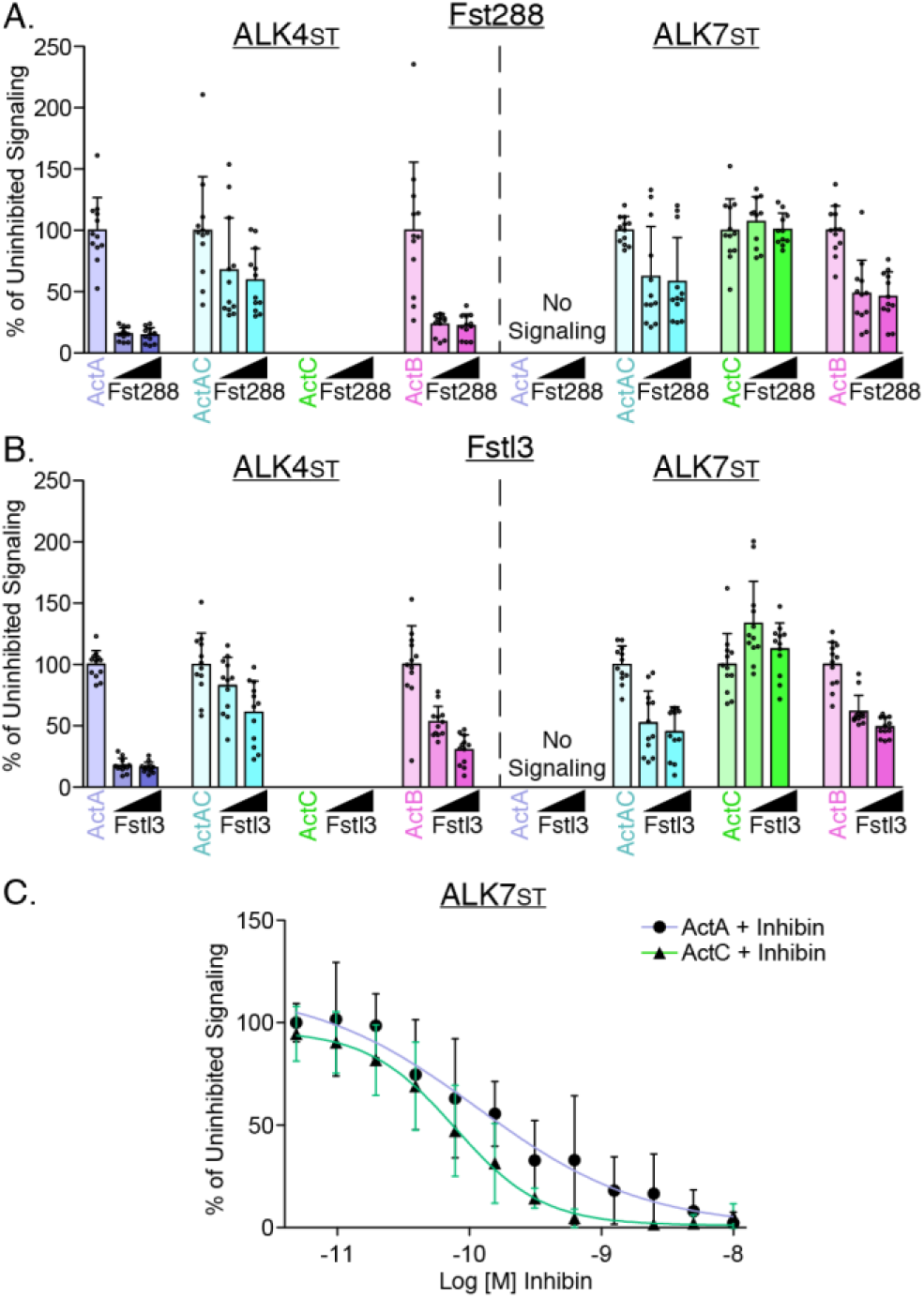
Activin C is resistant to inhibition by follistatin, but not inhibin A. (*A*) HEK293T (CAGA)_12_-luc cells were transfected with ALK4_st_ and ALK7_st_ and treated with SB-431542 and ActA, ActAC, ActC, or ActB (0.62nM) with increasing quantities (12.5 nM or 25 nM) of either Fst288 (*A*) or Fstl3 (*B*). (*C*) Luciferase assay following treatment with either ActA (ALK4_st_ signaling) or ActC (ALK7_st_ signaling) at a constant concentration (0.62 nM) along with titration of InhA. Each data point represents technical replicates within triplicate experiments measuring relative luminescence units (RLU) with bars displaying the mean ± SD. Data are represented as % of uninhibited.

Inhibins are ligand-like antagonists of activins that are formed from the heterodimerization of an InhβA or InhβB chain and the Inhα chain, resulting in the heterodimers inhibin A and B^41^. These heterodimers acts as a signaling dead molecules by binding type II receptors in a nonproductive receptor complex. To test whether inhibin A can antagonize ActC signaling, we titrated recombinant inhibin A against ActC in the above-described ALK7_st_ luciferase assay. Like ActA, ActC signaling was dose-dependently attenuated by InhA with an IC50 value of 0.08 nM (SD +/- .04 nM) as compared to 0.2 nM for ActA (SD +/- .2 nM) (Fig. 4C).

### Modeling of the Activin C ligand

Given the low affinity of ActC for the activin type II receptors and the ligand’s resistance to follistatin inhibition, we next examined what molecular differences within the activin class ligands could account for variation in ligand-receptor or ligand-follistatin interactions. A trio of residues (Ile340, Ala341, and Pro342; IAP motif; ActA) at the ligand knuckle are utilized during both type II receptor and follistatin binding^33,38,39,42^. Sequence alignment across the activin class reveals conservation of this motif in each ligand of the activin family, except for ActC and ActE (Fig. 5A). During complex formation between ActA:ActRIIB, the IAP motif forms the core of interactions with a series of hydrophobic residues in ActRIIB (Tyr60, Trp78, Phe101) (Fig. 5B and C). Additionally, this interface is engaged by Fst288, highlighting that the IAP motif is utilized by both antagonists and receptors (Fig. 5D). The core interactions involving the IAP motif are consistent across other structures within the activin class, such as GDF8:Fst288, GDF11:ActRIIB, and GDF11:Fst288^16,19^. In comparison, ActC contains a glutamine residue in place of the central alanine residue of the IAP motif. Generating a model of ActC (swissmodel to ActA; PDB: 7OLY) and aligning it to ActA reveals that Gln267 of ActC would be sterically unfavorable for interactions with both ActRIIB and Fst288 (Fig. 5C and D). Thus, we hypothesized that Gln267 in ActC might contribute to the reduced interaction with both the type II receptors and Fst288.

**Figure 5.**
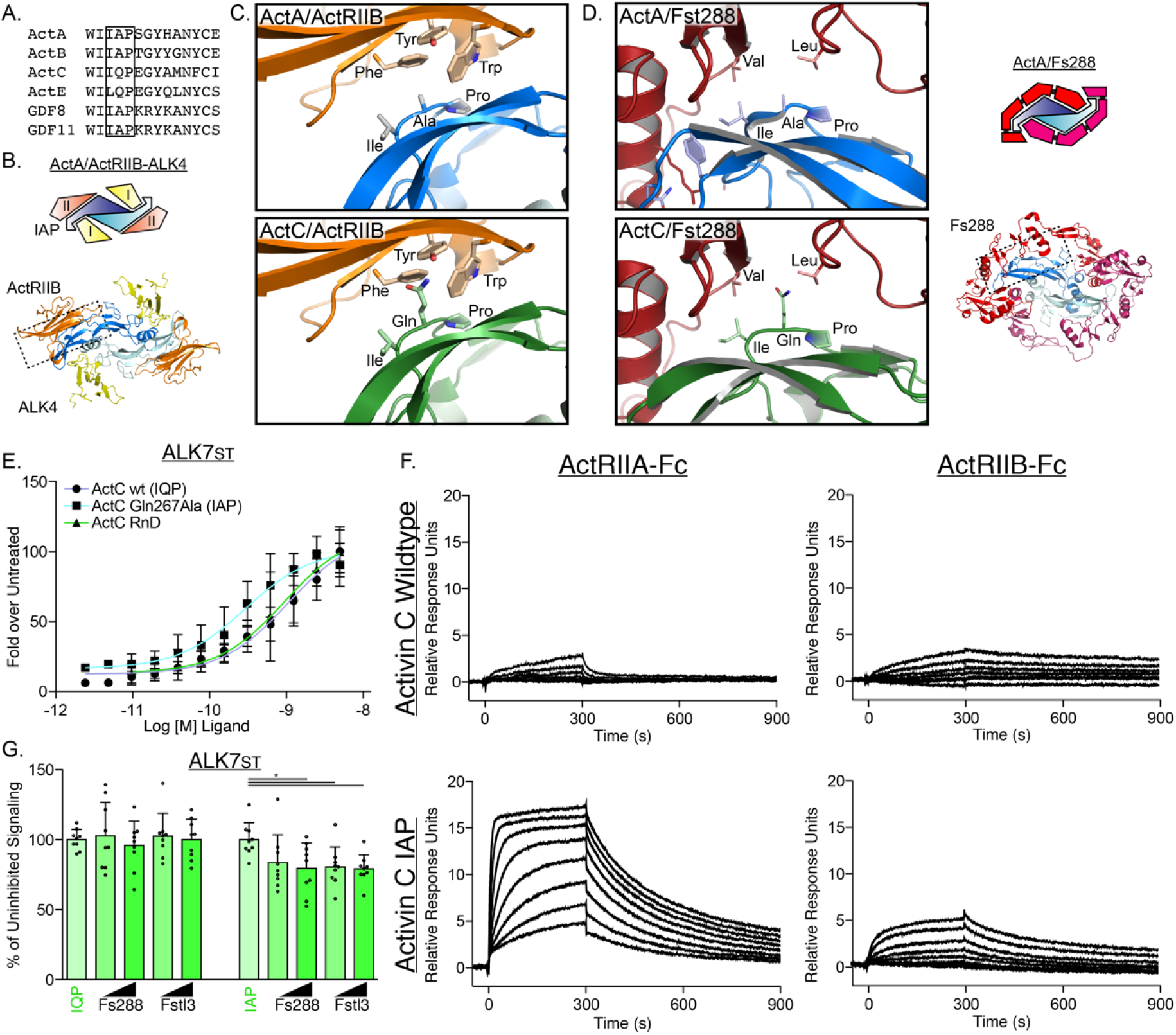
The Type II Interface of ActC is distinct from other activins. (*A*) Sequence alignment across the activin class displays critical differences at the canonical type II receptor binding site. IAP motif is boxed in *black*. (*B*) Structure and schematic representation of the ActA/ActRIIB/ALK4 complex (PDB: 7OLY). ALK4 is in *yellow*, ActRIIB is in *orange*, ActA is in *blue*. (C) Comparison of type II receptor interface between ActA/ActRIIB and the ActC/ActRIIB model, centered on the IAP (ActA) and IQP (ActC) motifs. The ActC model was built from (PDB: 7OLY)^38^. (*D*) Comparison (*left*) of the Fst288 interface between ActA/Fst288 (PDB: 2BOU) and the ActC/Fst288 model, centered on the IAP (ActA) and IQP (ActC) motifs. Schematic of ActA/Fst288 included (*right*) for clarification. (*E*) ALK7_st_-dependent luciferase assay following treatment with ActC purchased from RnD Systems, or recombinant ActC wildtype (IQP) or ActC Gln267Ala (IAP) transiently produced in HEK293T cells. Each data point represents the mean ± SD of triplicate experiments measuring relative luminescence units (RLU). (*F*) Averaged SPR sensorgrams of ActC wt (IQP) and ActC Gln267Ala (IAP) binding to protein-A captured ActRIIA-Fc or ActRIIB-Fc. Sensorgrams (*black lines*) are overlaid with fits to a 1:1 binding model with mass transport limitations (*red lines*). Each experiment was performed in duplicate. (*G*) ALK7_st_-dependent luciferase reporter assay following treatment of ActC wt (IQP) and ActC Gln267Ala (IAP) (0.62 nM) with increasing concentrations (12.5 nM or 25 nM) of either Fst288 or Fstl3. Each data point represents technical replicates within triplicate experiments measuring relative luminescence units (RLU) with bars displaying the mean ± SD.

To test this idea, we expressed and purified both recombinant wildtype (IQP) and Q267A (IAP) ActC from HEK293T cells. The IAP mutant had similar activity to the IQP wildtype form of ActC in the ALK7_st_-dependent CAGA-luc assay (Fig. 5E). As determined by SPR, ActC wildtype had low affinity for both ActRIIA-Fc and ActRIIB-Fc (Fig. 5F), consistent with binding data using recombinant ActC from R&D Systems. Replacement of the glutamine with alanine in ActC resulted in an increase in type II receptor binding, especially for ActRIIA-Fc (Fig. 5F). Next, we challenged ActC^Q267A^ with follistatin in the CAGA-luc assay. Here, ActC^Q267A^ was more inhibited by both Fst288 and Fstl3 than wildtype ActC (Fig. 5G). Taken together, these data support the hypothesis that the glutamine substitution in ActC relative to the other ligands of the activin class (ActA, ActB, GDF8 and GDF11) weakens the affinity for both the type II receptors and follistatin.

### Activin C signals similarly to activin B in mature adipocytes

Since we have shown that ActC can activate ALK7 in a cell-based luciferase assay, we next sought to determine the ligand’s capacity to signal via endogenous ALK7 in a biologically relevant cell type, adipocytes. To this end, we utilized both the preadipocyte cell line, 3T3-L1, and mature adipocytes derived from the stromal vascular fraction (SVF) of murine adipose tissue. Cells were differentiated over 4 days and maintained for 6 further days, where cell morphology and lipid droplets visibly increased, indicative of mature adipocytes (Fig. 6A). ActC stimulated SMAD2 phosphorylation (pSMAD2) in adipocytes differentiated from SVF but not 3T3-L1 cells (Fig. 6B and 6C). Notably, ALK7 (product of the *Acvr1c* gene) expression was significantly higher in SVF-relative to 3T3-L1-derived adipocytes (Fig. 6D). ActB stimulated pSMAD2 in both cell types, presumably via ALK4 in 3T3-L1 or a combination of ALK4 and ALK7 in differentiated adipocytes (Fig. 6B and 6C). In the mature (SVF-derived) adipocytes, both ActB and ActC induced pSMAD2 in a similar manner; however, Fst288 only blocked ActB action (Fig. 6C), consistent with the results above in the ALK7_st_-dependent luciferase assay (Fig. 4A). The neutralizing ALK7 antibody blocked ActC-induced pSMAD2 in mature adipocytes supporting that signaling was dependent on the ALK7 receptor (Fig. 6E). Interestingly, ActB induced pSMAD2 was only partly blocked in this assay, likely due to residual signaling via ALK4 (Fig. 6E). These results demonstrate that ActC is an ALK7-dependent signaling ligand and is follistatin resistant in a physiologically relevant context (Fig. 6C and E).

**Figure 6.**
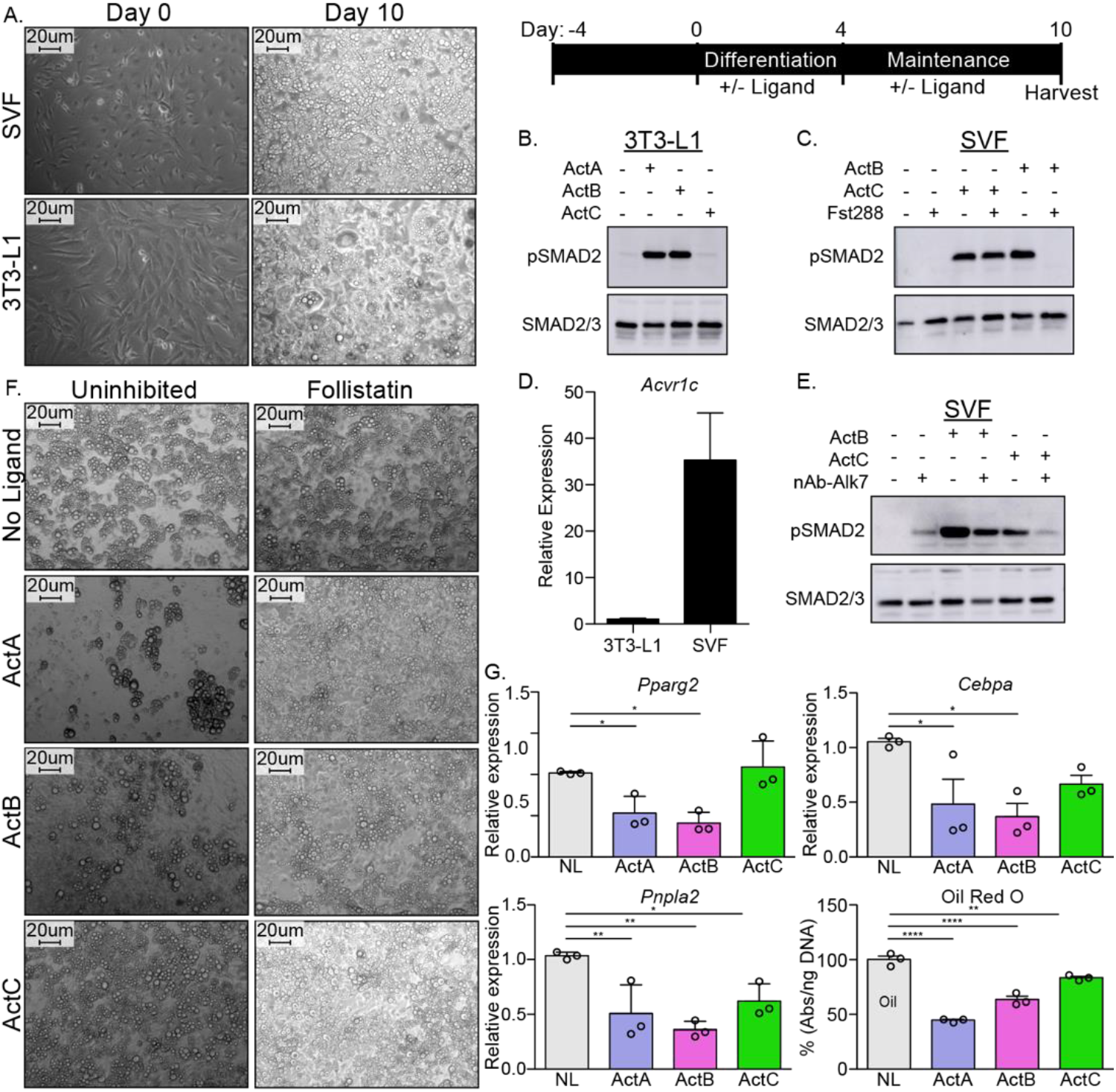
ActC activates SMAD2 through ALK7 in differentiated adipocytes. (*A*) Representative images of isolated adipose-derived stromal vascular fraction (SVF) or cultured 3T3-L1 cells prior to differentiation (*left*, Day 0) and following differentiation (*right*, Day 10). Scale bars are 20 μm. Schematic shown in *upper right* for visualization of timeline. (*B*) Western blot (WB) showing phosphorylated SMAD2 (pSMAD2) and total SMAD2/3 in 3T3-L1-derived adipocytes following treatment with ActA, ActB, or ActC (2 nM) for 1h. (*C*) WB following treatment of SVF-derived adipocytes with ActB or ActC (2 nM) with or without Fst288 (800 ng/ml) for 1h. (*D*) Quantitative PCR (RT-qPCR) of *Acvr1c* expression in differentiated 3T3-L1 cells and SVF adipocytes. Bars display mean ± SD of three experimental replicates. (*E*) WB following treatment of SVF-derived adipocytes with ActB or ActC (2 nM) in the presence or absence of a neutralizing antibody targeting ALK7 for 1h. (*F*) Representative images of SVF-derived adipocytes following treatment with ActA, ActB, or ActC during differentiation with or without Fst288. (*G*) RT-qPCR of target genes *Pparg2, Cebpa*, and *Pnpla2* following treatment with ActA, ActB, or ActC during differentiation. Oil Red O quantification based on images in (*F*). Significance is represented as: * p<0.05, ** is p<0.01, *** p<0.001 and **** p<0.0001. Each experiment was performed in triplicate. While representative westerns are shown, supplemental westerns can be found in SI appendix Fig. S3.

ActB has dual effects on adipogenesis, and its function depends on the relative expression of ALK4 and ALK7 during the process of adipocyte commitment and differentiation^24,29,43,44^. ActA or ActB exposure during differentiation of SVF cells, when ALK7 levels are low, inhibits adipogenesis (Fig. 6F). Treatment with ActC at this early stage did not affect adipogenesis (Fig. 6F). Follistatin antagonized the anti-adipogenic effects of both ActA and ActB, restoring normal adipogenesis and lipid droplet formation (Fig. 6F). Furthermore, gene expression of both *Pparg2* and *Cebpa*, essential transcription factors for adipogenesis, was impaired by ActA or ActB, but not ActC (Fig. 6G). However, ActC significantly reduced both *Pnpla2* expression and lipid content, consistent with the late-stage, proadipogenic effects of ActB-ALK7 signaling (Fig. 6G)^43,45-47^.

## Discussion

The binding/signaling profiles of some members of the activin class (ActA, ActB, GDF8, and GDF11) of TGFβ ligands have been largely characterized, where each member exhibits differential specificity for both the type II receptors, ActRIIA and ActRIIB, and the type I receptors: ALK4, ALK5, and ALK7. ActA is limited to a single type I receptor, ALK4, and has little type II receptor preference, while GDF11 can promiscuously signal through ALK4, ALK5, and ALK7 and seemingly favors interaction with ActRIIB^16^. The receptors for ActC have remained largely unknown in part due to the initial characterization of ActC as a non-signaling molecule^5^. In this study, we identified ActC as a bona fide signaling ligand with distinct molecular properties from other activin class ligands.

ActC signals through ALK7, whereas ActAC uses both ALK4 and ALK7. Thus, ligands that contain an InhβC subunit can bind and act through ALK7. This is similar to what was previously described for ligands containing an InhβB subunit, like activin B and activin AB. In contrast, ActA does not signal through ALK7. Interestingly, the heterodimer ActAC was more potent than ActC, which has similarly been reported for other heterodimers in the family, such as BMP2/4 and BMP2/7^48,49^. This might be a results of different type I receptor binding epitopes that are formed in the heterodimer versus the homodimer.

The molecular basis for type I receptor specificity remains an intriguing aspect of the evolution of the activin class ligands. Initial studies implicated the wrist region, including the prehelix loop as a major contributor towards type I receptor specificity, as swapping this region could alter type I receptor specificity^35^. More recent studies have identified residues in the fingertip region of the ligand as also have a major role in type I receptor specificity^16,50^. Given the low affinity nature of the type I receptors for ligands across the activin family, the current thought is that subtle differences at the type I:ligand interface dictate receptor specificity. Interestingly, fingertip residues that are important for ActA and GDF11 binding to type I receptors are divergent in ActC (Fig. S4) and could account for the latter’s lack of signaling through ALK4. Most notably, a recent crystal structure of ActA in complex with ALK4 shows that D406 of ActA forms a hydrogen bond with the mainchain of ALK4. The corresponding residue is an arginine in ActC^20^. On the receptor side, the β4-β5 loop is important for ligand recognition and is shorter in ALK7 than ALK4 and ALK5, possibly to accommodate the larger arginine residue, which will extend towards this loop. Certainly, structures of ALK7 in complex with ActC or other ligands will help determine how specificity for ALK7 is acquired. Regardless, it seems that differences in the ligand fingertip and/or prehelix loop, coupled with differences in the receptor β4-β5 loop, dictate type I receptor specificity in the activin class.

In general, activin class ligands bind activin type II receptors with high affinity^16,20,33,42^. Conversely, ActC exhibits weak binding to both ActRIIA and ActRIIB. ActAC binds the type II receptors, but with reduced affinity compared to ActA. Thus, the InhβC subunit appears to reduce affinity for the activin type II receptors. Nevertheless, the type II receptors are required for ActC signaling, as the ligand’s activity was abrogated by activin type II receptor neutralizing antibodies and inhibin A. Unlike the other members of the activin class, ActC binds both type I and type II receptors with low affinity but is still able to signal. One possible explanation is that ActC binds to the type I and type II receptors cooperatively. This is supported by SPR studies with the hetero-receptor combination of ALK7-ActRIIB, which has a higher affinity for ActC than either receptor alone (Fig. 2). It has been suggested that exogenous ActC can directly antagonize ActA signaling^8^. One proposed mechanism is that ActC would be a competitive inhibitor towards the ActA receptors. Our data challenges this idea as ActC binds weakly to ActRIIA, ActRIIB, and the fusion ActRIIB-ALK4-Fc. Thus, the main mechanism of ActC antagonism of ActA is likely through heterodimer formation, where the ActAC ligand signals less potently through ALK4, while gaining the ability to signal through ALK7, when present.

Another unexpected characteristic of ActC is its interaction or lack thereof with the extracellular antagonists, the follistatins. Given ActC’s structural similarity to other activin class members, it was unexpected that neither Fst288 nor Fstl3 inhibited ActC. Similarly, suppression of ActAC signaling was less significant compared to the other activin ligands ActA and ActB (Fig. 4). This indicates that the inhβC chain limits follistatin binding and confers resistance to antagonism. The biological implications of follistatin resistance will need to be further explored, but it is tempting to speculate that the presence of follistatin would interfere with ligands that signal through ALK4, providing a permissive environment for the lower affinity ActC to bind type II receptors and signal via ALK7. Additionally, the presence of follistatin, while limiting ActA and ActB signaling, would still permit ActAC signaling via ALK4.

Having low affinity for the type II receptors and resistance to follistatin distinguishes ActC from the other members of the activin class and raises the question as to what confers differences in ligand properties, especially given their >50% sequence identity. Comparison across the activins revealed conservation of the shared type II/follistatin binding surface, except for a single residue, centrally located in the interface. While most activins have an alanine at this position, InhβC contains a glutamine akin to the glutamate within the TGF-β’s (TGF-β 1-3) (Fig. S4). Converting this residue to an alanine in ActC resulted in a ligand with higher affinity for ActRIIA and increased sensitivity to follistatin. While not the only molecular difference, it appears that InhβC has evolved a single substitution centrally located in a major binding epitope relative other activin class members that suppresses its interaction with follistatin and type II receptors. Comparison across different species shows this deviation is conserved in mammals (Fig. S5). Interestingly, fish are divergent and possess the alanine version of InhβC similar to InhβA and InhβB. This bifurcation in conservation suggests differentially evolved activin ligands exist in the two taxa and might provide clues as to the biological function of ActC in different species.

Physiological roles for ActC and ActAC have not yet been established. However, given the ability of the ligands to signal via ALK7, we turned our attention to adipocytes. Human adipose tissue is a major site of both ActB and ALK7 expression, where the pair induces pro-obesity signaling outcomes, such as catecholamine resistance or inhibition of lipolysis^29,43,51^. In this study, we show that ActC regulates adipocyte differentiation differently than ActA or ActB. While both ActA and ActB exhibit potent inhibitory effects in cultured adipocytes, ActC does not. This difference is explained, in part, by the ability of ActA and ActB, but not ActC, to signal via ALK4, which is present in pre-adipocytes and throughout their maturation and differentiation. In contrast, ActC can signal in adipose tissue only once ALK7 is expressed, which occurs in the later stages of adipocyte differentiation. In this context, ActC negatively regulates lipase expression, lipid content, and elicits a SMAD2 response similar to that of ActB. In addition to ActB and ActC, the TGFβ ligand, GDF3, can also signal through ALK7. GDF3 signaling is supported by the co-receptor Cripto and is implicated in the regulation of energy homeostasis and adipocyte function^27^. Thus, taken together, it appears that several TGF-β ligands have evolved the ability to signal in adipocytes depending on the receptor/co-receptor profile.

Given that ActC expression and secretion is highest within the liver, a tissue with low ALK7 expression, it is possible that ActC acts as a hepatokine functioning systemically in fat regulation^51,52^. Unlike ActB, ActC is not antagonized by follistatin and may therefore serve as an uninhibited, basal signal in the face of variable follistatin expression, such as during thermogenesis in adipocytes following cold exposure^53^. Another possible role might be during liver regeneration, as ActAC and ActC have been observed in serum, coinciding with a surge of follistatin expression^54^.

Another member of the activin class, ActE, is expressed highest in the liver and has been implicated as a hepatokine with a role in energy homeostasis; however, whether ActE can signal and, if so, through which receptors, has yet to be determined^55^. Interestingly, ActE has a glutamine in position 267, similar to ActC, and a leucine in position 265, replacing the conserved isoleucine (Fig. S4). These amino acid differences likely reduce ActE’s affinity for the type II receptors and follistatin similar to ActC. Given these similarities it is possible that there is functional overlap between ActC and ActE, possibly in the liver-adipose signaling axis.

Throughout the body, there are a variety of activins with differential receptor and antagonist binding, yielding a variety of potential signaling capacities (Fig. 7). Our results indicate that ActC can act as a canonical TGFβ ligand, transducing SMAD2/3 responses similar to ActA and ActB, while avoiding inhibition through the follistatin family of antagonists. This observation challenges current thinking that ActC acts solely as an activin antagonist. Different than ActB, which can signal through both ALK4 and ALK7, ActC is specific for ALK7 and shows a preference for the type II receptor ActRIIA. Future studies will need to address the different biological roles of ActC and ActAC signaling through ALK7, with an initial focus on adipose tissue.

**Figure 7.**
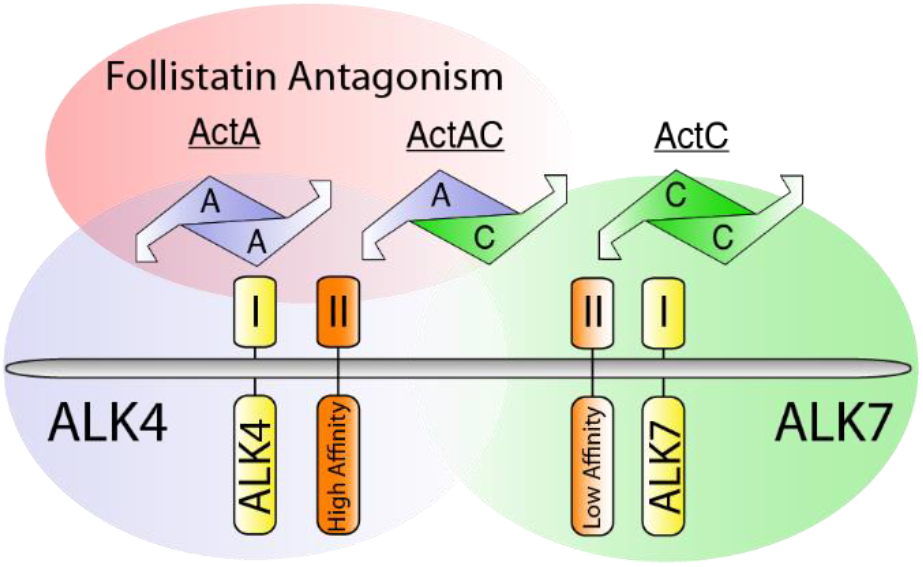
Differences between ActA and ActC in type I receptor specificity, type II receptor affinity, and follistatin antagonism. Gradients of follistatin antagonism (*red*), ALK4-dependent signaling (*blue*), ALK7-dependent signaling (*green*), and activin type II receptor affinity (*orange*) for ActA, ActAC, and ActC. Ligands and type I receptors are shown schematically.

## Author Contributions

Each author contributed to research design; E.J.G., L.O., and E.K., and E.B. performed research; R.C. and R.K. contributed reagents/analytic tools; E.J.G and T.B.T wrote the manuscript. D.J.B., L.O. R.C. and E.K., edited the manuscript.

---

**Supplemental Figure 1.**
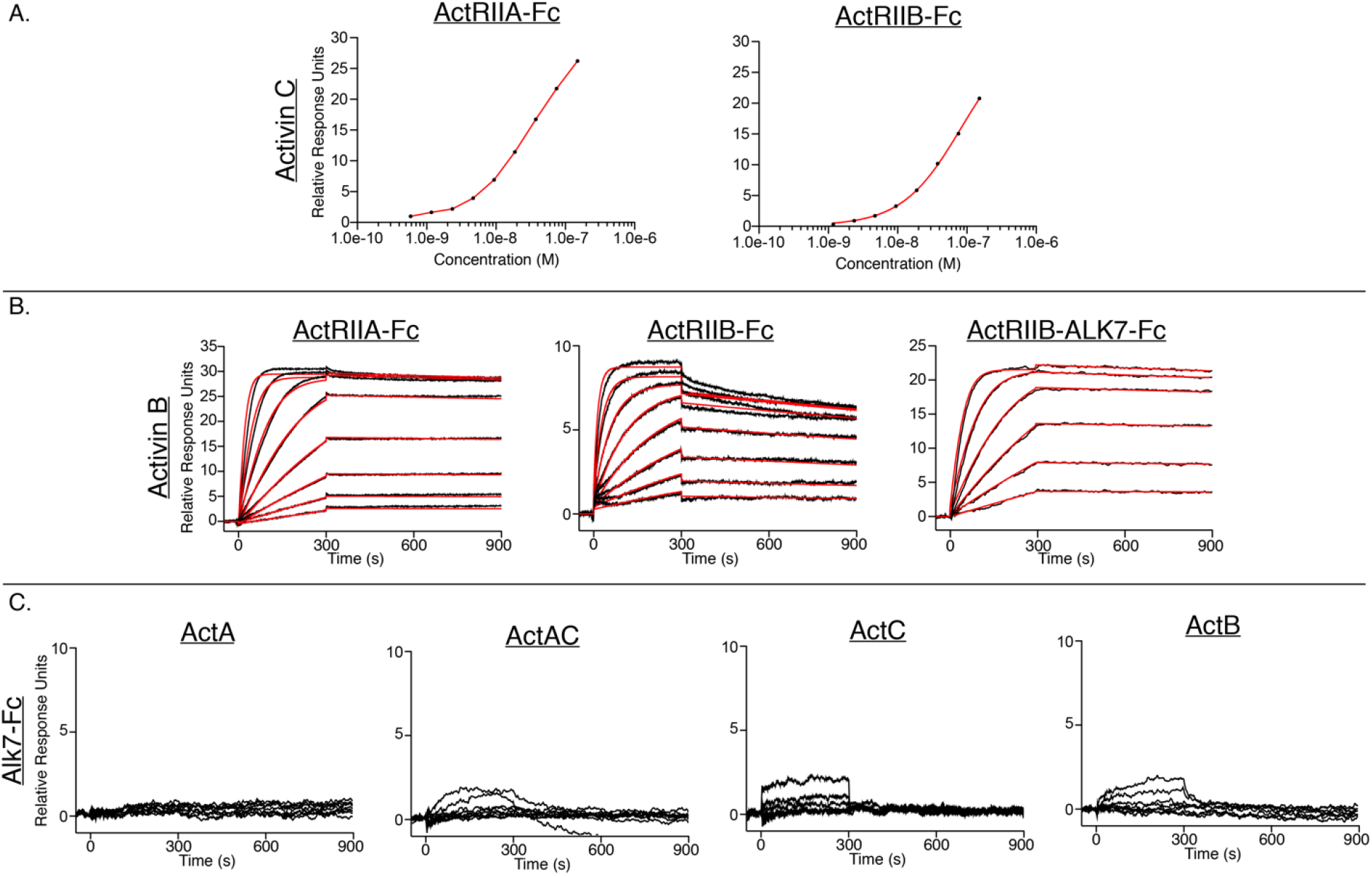
Additional surface plasmon resonance sensorgrams of ActA, ActAC and ActC binding to Alk7 and ActB binding to different receptors. (A) Steady state analysis of the following experiments in Fig. 2: ActC binding to captured ActRIIA-Fc and ActRIIB-Fc. (B) Representative SPR sensorgrams of ActB binding to protein-A captured ActRIIA-Fc, ActRIIB-Fc, or ActRIIB-ALK7-Fc. Sensorgrams (*black lines*) are overlaid with fits to a 1:1 binding model with mass transport limitations (*red lines*). (C) Representative SPR sensorgrams of ActA, ActAC ActC and ActB with Alk7-Fc captured on a protein A chip. Each experiment was performed in duplicate with the kinetic parameters, if available, summarized in SI appendix Table S1.

**Supplemental Table 1.** SPR Kinetic Analysis.

| Analyte | Ligand | $k_a$ (M<sup>-1</sup>s<sup>-1</sup>) x 10<sup>6</sup> | $k_d$ (s<sup>-1</sup>) x 10<sup>-4</sup> | $K_D$ (pM)<sup>a</sup> |
| --- | --- | --- | --- | --- |
| Activin A | ActRIIA-Fc | 7.2 ± 1.4 | 1.6 ± 0.33 | 22 ± 0.030 |
| Activin A | ActRIIB-Fc | 7.7 ± 1.0 | .60 ± .055 | 8.1 ± 1.8 |
| Activin A | ActRIIB-ALK4-Fc | 7.0 ± 0.77 | 2.1 ± 0.20 | 30 ± 6.2 |
| Activin A | ActRIIB-ALK7-Fc | 2.9 ± 0.15 | 8.9 ± 0.09 | 310 ± 12 |
| Activin AC | ActRIIA-Fc | 2.9 ± 0.040 | 4.3 ± 0.39 | 150 ± 12 |
| Activin AC | ActRIIB-Fc | 2.4 ± 0.16 | 2.1 ± 0.080 | 90 ± 2.6 |
| Activin AC | ActRIIB-ALK4-Fc | 2.1 ± 0.34 | 9.2 ± 0.26 | 460 ± 88 |
| Activin AC | ActRIIB-ALK7-Fc | 5.6 ± 0.23 | 2.8 ± 0.12 | 51 ± 0.050 |
| Activin C | ActRIIA-Fc | Transient Binding | Transient Binding | - |
| Activin C | ActRIIB-Fc | Transient Binding | Transient Binding | - |
| Activin C | ActRIIB-ALK7-Fc | .25 ± .0075 | 5.7 ± 0.060 | 2200 ± 44 |
| Activin B | ActRIIA-Fc | 7.7 ± 0.15 | .73 ± 0.0020 | 9.5 ± 0.21 |
| Activin B | ActRIIB-Fc | 7.2 ± 0.22 | 2.1 ± 0.36 | 30. ± 5.9 |
| Activin B | ActRIIB-ALK7-Fc | 14 ± 0.0 | .58 ± 0.10 | 4.0 ± 0.70 |

<sup>a</sup>All kinetic parameters were analyzed using the Biacore T200 evaluation software using a 1:1 binding model and are the average of two independent, replicate experiments.

**Supplemental Figure 2.**
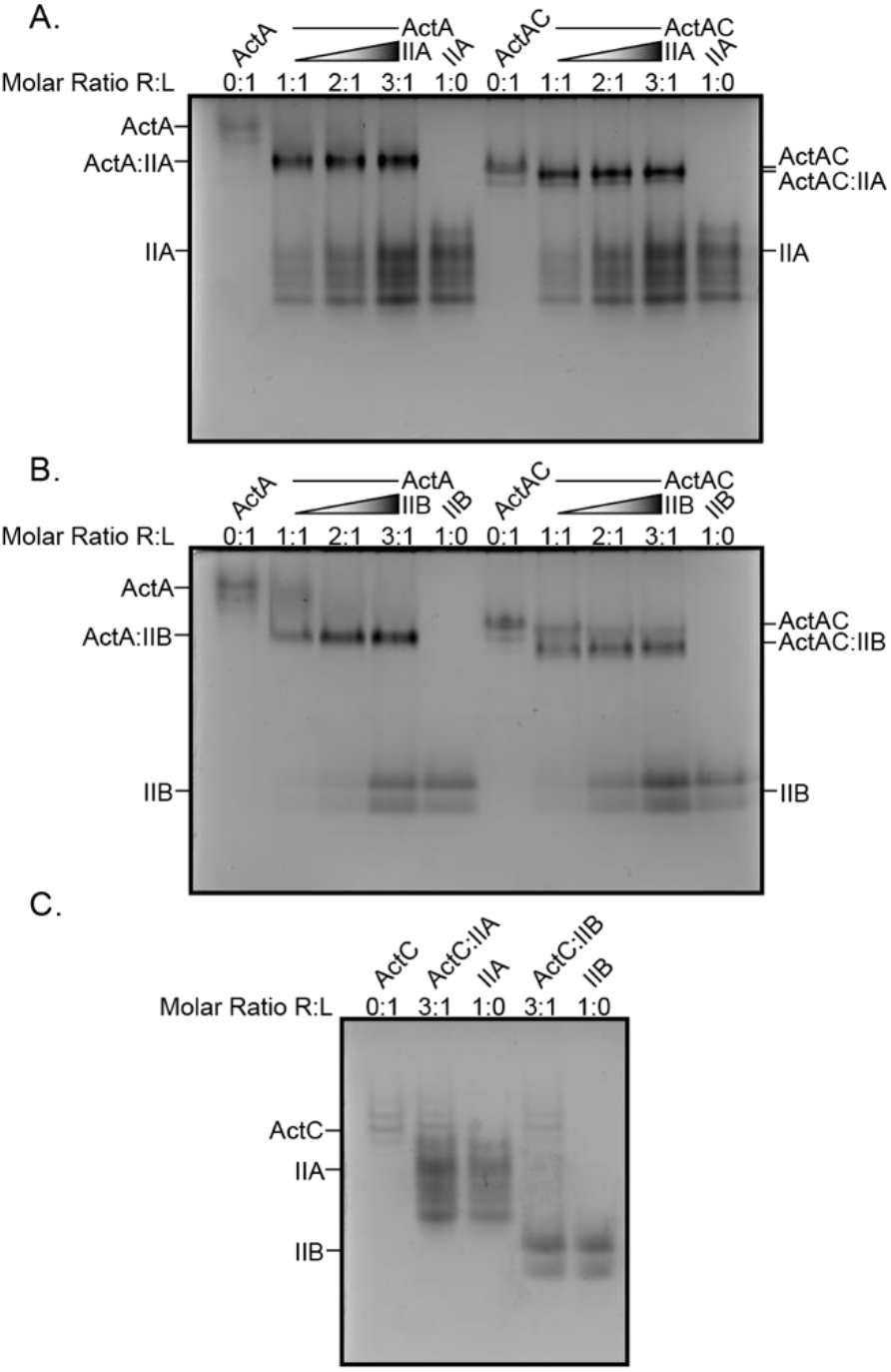
Native gel analysis of type II receptors and ActA, ActAC, and ActC. Native PAGE analysis of ActRIIA (*A*) and ActRIIB (*B*) with ActA and ActAC. Binary complexes were formed by titrating receptor from 1:1 to 3:1 molar ratio against constant ligand. (*C*) Native PAGE analysis of 1:3 receptor:ligand molar ratio with ActC.

**Supplemental Figure 3.**
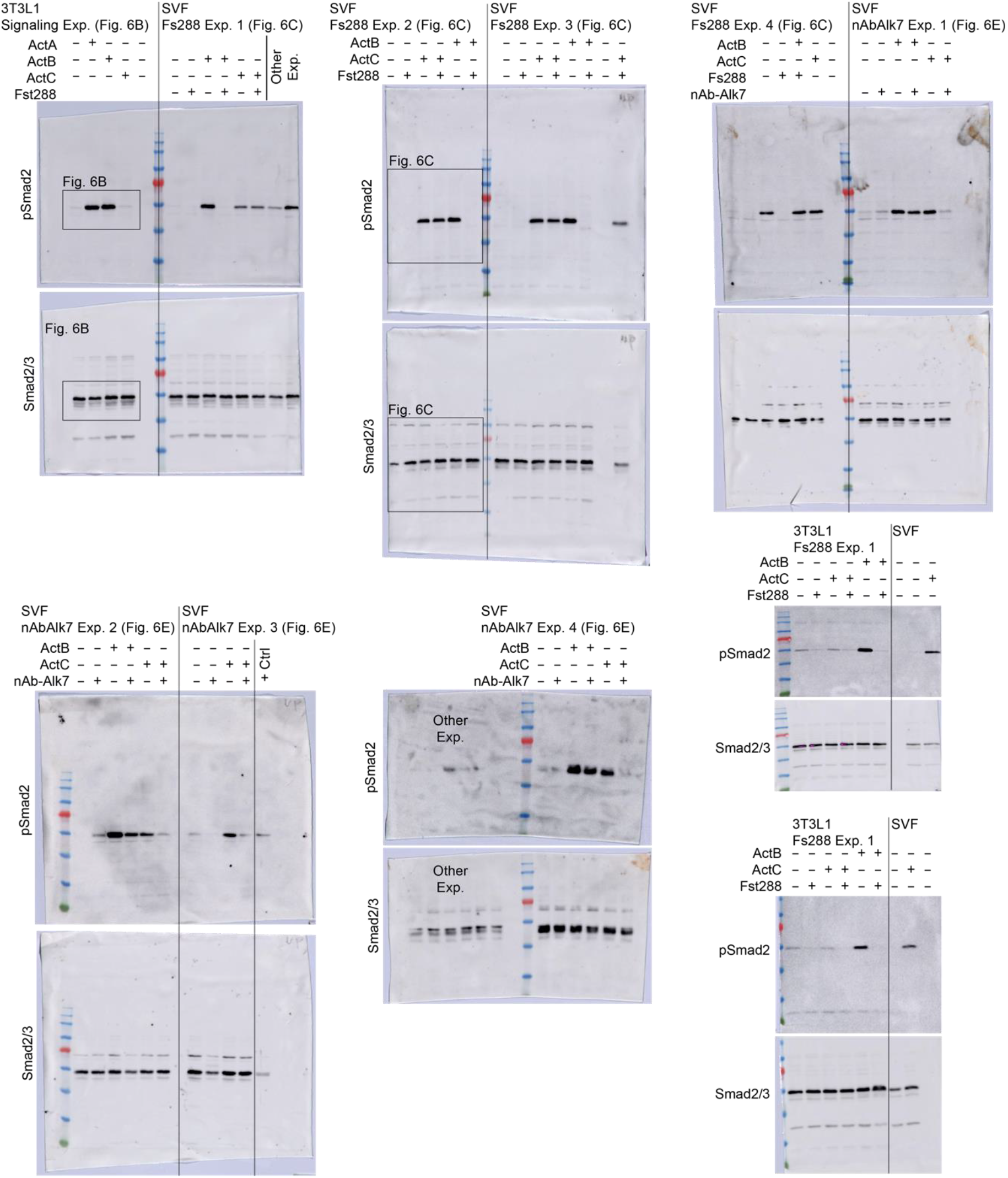
Supplemental Adipocyte-pSMAD2/SMAD2/3 western blots. Supplemental westerns for representative blots shown in Fig. 6. Boxes are drawn to display which data were used for figure generation. Antibodies used: pSMAD2 (Cell Signaling, 138D4) and SMAD2/3 (Millipore, 07-408).

**Supplemental Figure 4.**
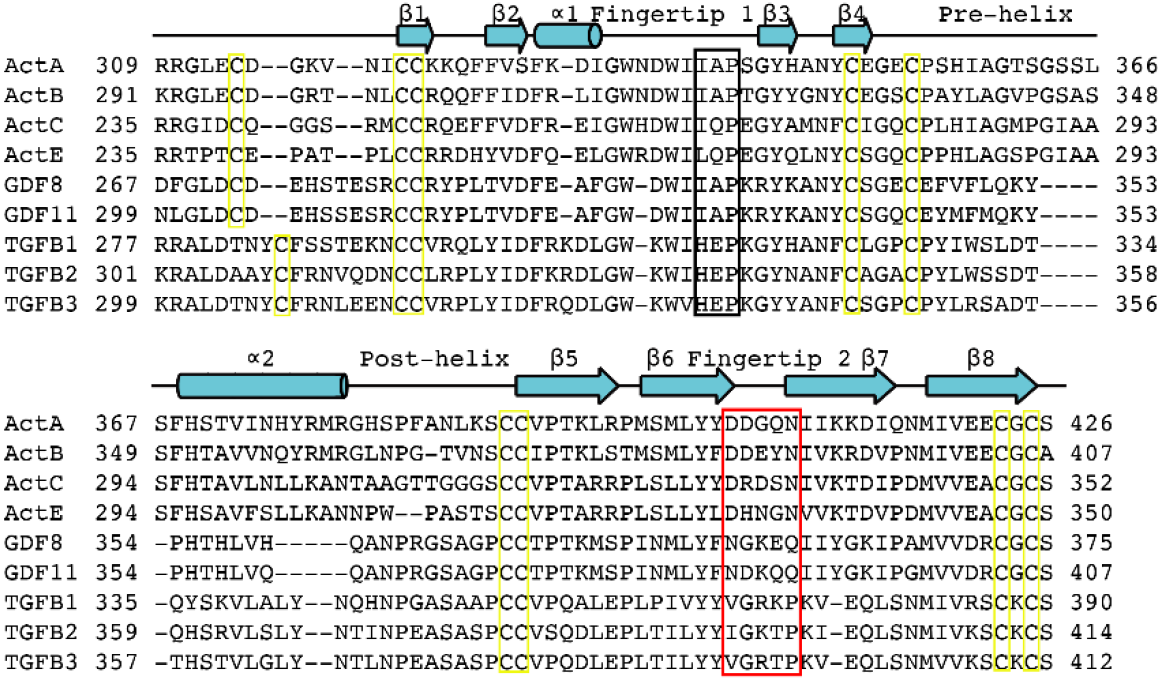
Sequence alignment of Activin and TGFβ ligands. Sequence comparison of the mature ligand for human Activin subclass (ActA, ActB, ActC, ActE, GDF8 and GDF11), TGFβ1, TGFβ2, and TGFβ3. Numbering includes the signal sequence and prodomain (not shown). Fingertip residues are boxed in red. The conserved IAP motif is highlighted with a black box. Secondary structure elements are represented as arrows or cylinders for β-strands and α-helices, respectively. Disulfide bonds are boxed in yellow.

**Supplemental Figure 5.**
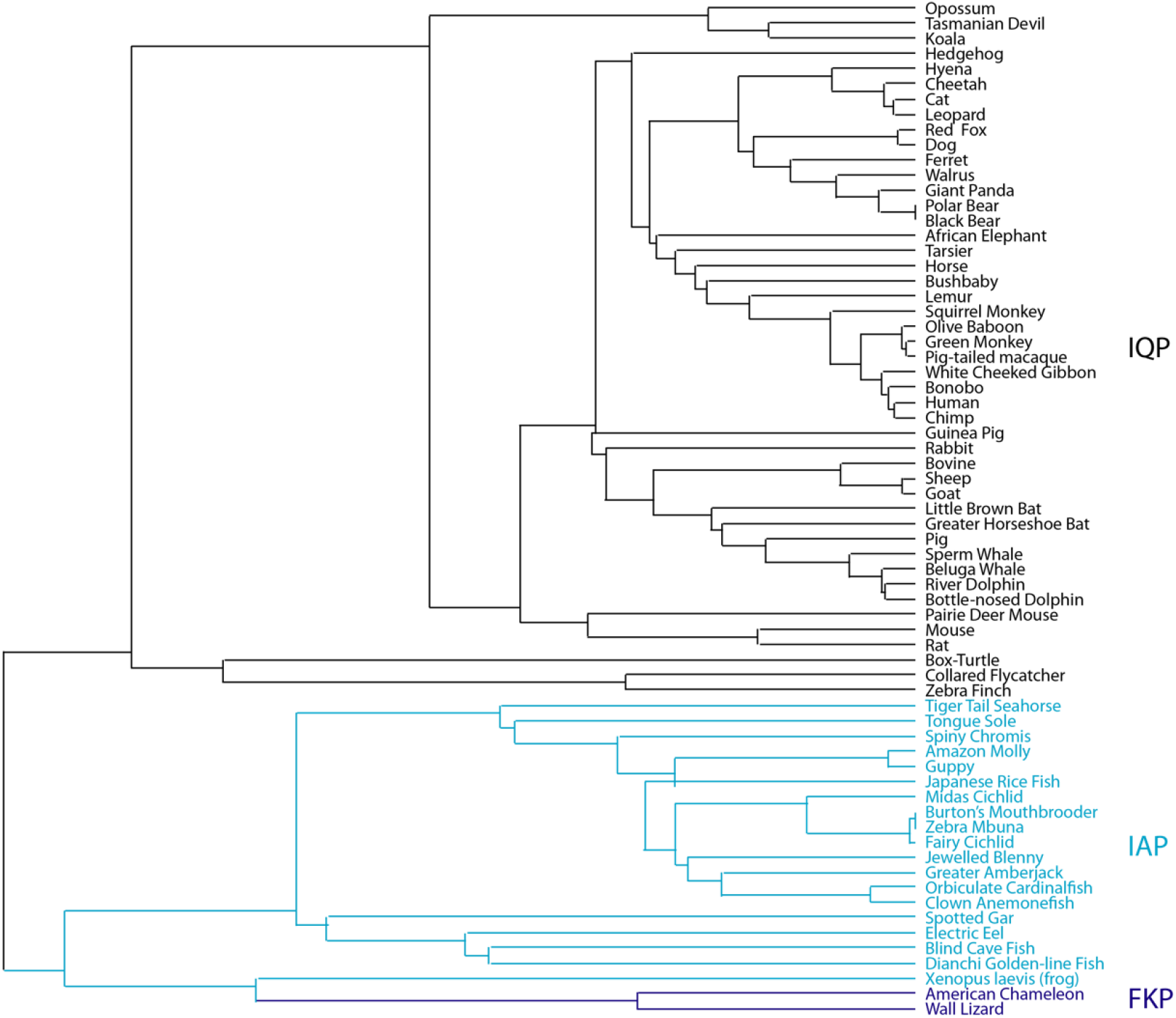
The Phylogenetic History of ActC. Phylogenetic analysis of full-length ActC across a large variety of species with focus on the type II receptor interface variance. Species with the IQP variant shown in *black*, with the IAP variant in *blue*, and with the FKP variant in *purple*.

**Supplemental Table 2.**
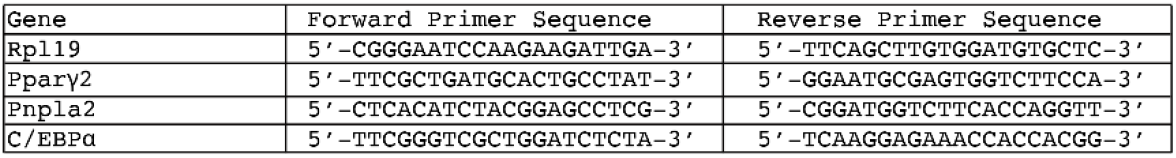
qPCR primer sequences.

